# Dopamine dynamics in human anterior cingulate cortex during Pavlovian-instrumental conflict

**DOI:** 10.64898/2026.04.02.716040

**Authors:** Azadeh Nazemorroaya, Seth R. Batten, Itamar Grunfeld, Alexis Torres, Xavier Celaya, Olivia Moreland, Chloe Lattuca, Ava Wagle, Devin Nikjou, Leonardo S. Barbosa, Terry Lohrenz, Pearl Chiu, Gene A. Brewer, Samuel M. McClure, Mark R. Witcher, Robert W. Bina, P. Read Montague, Peter Dayan, Dan Bang

## Abstract

Dopamine is believed to modulate not only instrumental learning about the link between states, actions, and outcomes but also reflexive behaviours, such as a Pavlovian bias to approach in rewarding states and freeze in aversive ones. We studied these dual roles in the human brain, by combining intracranial dopamine recordings from the anterior cingulate cortex (ACC)— a region implicated in behavioural and cognitive control — with a motivational Go/NoGo task involving conflict between instrumental and Pavlovian action selection. We found evidence that dopamine in the ACC is involved in evaluating whether Pavlovian responding should guide behaviour. This computational motif was observed across multiple task events, including in response to rewards and punishments, and in analyses based on a reinforcement learning model. Our results indicate that dopamine supports learning at the more abstract level of behavioural policies in addition to the more concrete levels of states and actions.

## 1 Introduction

Dopamine (DA) — one of the brain’s major neuromodulators — is crucial for healthy function. The DA system, which originates in small midbrain nuclei and projects widely throughout the brain (Trudeau and Denis, 2025), is believed to support fundamental cognitive and behavioural processes, including motivation, learning and movement (Berke, 2018; Bromberg-Martin et al., 2010; Wise, 2004). Consistent with this broad range of functions, disturbances in DA signalling have been implicated in psychiatric and neurological disorders that affect millions of people worldwide, such as schizophrenia (Leucht et al., 2025), substance use disorders (Koob and Volkow, 2016) and Parkinson’s disease (Poewe et al., 2017). Similarly, pharmacological treatments that target the DA system can modulate cognition and behaviour. For example, while being able to alleviate the effects of the loss of DA neurons on movement in Parkinson’s disease, DA replacement therapy can induce impulse control disorder (Voon et al., 2017).

DA is likely to support cognition and behaviour in multiple ways, given the diversity of cell types, receptor types and projection targets (Awatramani and Poulin, 2025; Trudeau and Denis, 2025; Watabe-Uchida and Amo, 2025). One recurring computational motif is an involvement in aspects of value (Schultz, 2025; Watabe-Uchida Mitsuko, 2025). Most prominently, a large body of research indicates that a central role of DA is to signal errors in the prediction of reward, as conceived in the artificial intelligence field of reinforcement learning (RL; Sutton and Barto, 2018). In their simplest form, reward prediction errors (RPEs) can be defined as the difference between received and expected levels of reward. RPEs provide a teaching signal for learning the value of states and actions in the world, as often studied in forms of instrumental or operant conditioning. Neural data that conform to the RPE theory of DA has been observed in animals (Cohen et al., 2012; Glimcher, 2011; Hart et al., 2014; Montague et al., 1996; Schultz et al., 1997) and humans (Batten et al., 2024; Kishida et al., 2016; Moran et al., 2018; Sands et al., 2023) — alongside recent evidence that DA also reports hypothetical RPEs, such as RPEs over the value of actions not taken (Kishida et al., 2016; Moran et al., 2018), a counterfactual teaching signal that can further speed up learning (Bennett et al., 2022).

However, DA appears not only to modulate estimates of the value of states and actions but also to be involved in the realisation of reflexive behavioural tendencies (Collins and Frank, 2014; Corbit and Balleine, 2015; Gentry et al., 2016; Jaskir and Frank, 2023; Lex and Hauber, 2008; Syed et al., 2016). Perhaps the best-known example of such tendencies is Pavlovian responding, whereby agents tend to activate behaviour in states that predict reward and inhibit behaviour in states that predict punishment (Guitart-Masip et al., 2014). When the demands of instrumental and Pavlovian responding are congruent — when an agent must emit an action (Go) to gain a reward or inhibit an action (NoGo) to avoid a punishment — there is no conflict for the agent. But when their demands are incongruent — when it is necessary to inhibit an action to gain a reward or to emit an action to avoid a punishment — a conflict arises. This conflict is well-established psychologically (Dayan et al., 2006) and is typically studied using motivational Go/NoGo tasks that manipulate the required action in a state (Go versus NoGo) and/or state valence (reward versus punishment) (Guitart-Masip et al., 2014). Largely based on evidence from the ventral striatum in rodents, DA has been implicated in aspects of Pavlovian-instrumental conflict (Cohen et al., 2012; Corbit and Balleine, 2015; Gentry et al., 2016; Hart et al., 2014; Lex and Hauber, 2008; Oleson et al., 2012; Syed et al., 2016), and indeed its resolution as an example of cognitive control (Lloyd and Dayan, 2023). However, perhaps due to its behavioural complexity, animal studies have not used a motivational Go/NoGo task that fully orthogonalises the required action in a state and state valence to cover all four scenarios. Human studies have combined variants of such a task with pharmacological manipulation of the DA system (Guitart-Masip, Chowdhury, et al., 2012; Guitart-Masip et al., 2014; Swart et al., 2017), but this approach does not address the role of DA at the fast timescales probed in animals.

Here, to understand how Pavlovian-instrumental conflict engages the DA system, we leveraged intracranial human electrochemistry (Bang et al., 2020; Batten et al., 2024, 2025; Kishida et al., 2016; Moran et al., 2018) to record sub-second DA dynamics in participants with epilepsy. During the recordings, participants performed a motivational Go/NoGo task which fully orthogonalises the required action in a state and state valence (Guitart-Masip, Huys, et al., 2012). We recorded from the anterior cingulate cortex (ACC), which receives substantial DA projections (Berger et al., 1991), can regulate the activity of DA neurons in the midbrain (Beier et al., 2015; Gariano and Groves, 1988), and has been implicated in multiple control functions, such as error detection, conflict monitoring, and the adaptation of behavioural strategies to environmental demands (Carter et al., 1998; Kerns et al., 2004; Kolling et al., 2016; MacDonald et al., 2000). In the context of Pavlovian-instrumental conflict, human fMRI and EEG indicates that the ACC biases striatal action selection in a valence-dependent manner (Algermissen et al., 2022, 2024). However, the role of DA at the level of the ACC is not well-understood, with computational accounts of DA largely based on recordings of midbrain DA neurons or DA dynamics in sub-cortical regions.

In the same way that DA release in the ventral striatum is believed to evaluate the extent to which Go responses are appropriate, we found evidence that DA concentrations in the ACC are involved in evaluating the extent to which Pavlovian responding is appropriate. In brief, DA signalled whether an action about to be made is Pavlovian congruent, whether erroneous initiated actions are surprising from the perspective of Pavlovian tendencies, and whether action outcomes favour Pavlovian responding. These results were observed in both model-agnostic analyses based on task-defined variables and model-dependent analyses informed by an RL model fitted to task behaviour. Our unique data from the human brain supports a hypothesis that DA is involved in RL at more abstract levels — here, the degree to which a default policy, Pavlovian responding, should guide behaviour — in addition to the more concrete levels of states and actions.

## 2 Results

### 2.1 Behavioural task and electrochemical recordings

Four participants with epilepsy performed an orthogonalised motivational Go/NoGo task (Guitart-Masip, Huys, et al., 2012) in an epilepsy monitoring unit (EMU). One participant performed the task twice on separate days, making for a total of five datasets. For convenience, we refer to each dataset as a participant/patient. In the task, participants had to learn by trial-and-error to press a button (Go) or withhold from pressing a button (NoGo) in order to increase the probability of winning a reward (Win) or avoiding a punishment (Avoid) (**Figure 1A**; see Methods for task details). In total, there are four experimental conditions, or states, denoted as: Go+ (Go-to-Win), Go- (Go-to-Avoid), NoGo+ (NoGo-to-Win) and NoGo- (NoGo-to-Avoid). By orthogonalising the required action in a state (Go/NoGo) and state valence (Win/Avoid), the task allows us to examine not only the separate contributions of action and value but also how they may interact to shape behavioural and neural responses. Specifically, the task manipulates the congruence, or conflict, between the appropriate cue-response mapping acquired through instrumental learning and Pavlovian biases to approach in rewarding contexts and avoid in aversive ones. In Pavlovian-congruent states (Go+ and NoGo-), instrumental and Pavlovian responding favour the same action. By contrast, in Pavlovian-incongruent states (Go- and NoGo+), instrumental and Pavlovian responding favour opposite actions. While participants performed the task, we obtained electrochemical estimates of DA levels from depth electrodes implanted into the ACC for the clinical purpose of epilepsy seizure monitoring (see **Figure 1B** for recording sites). Our electrochemical protocol, which has been validated in earlier work (Bang et al., 2023; Batten et al., 2025), provides ten DA estimates per second (see **Figure 1C** for evaluation and Methods for details on protocol).

**Figure 1.**
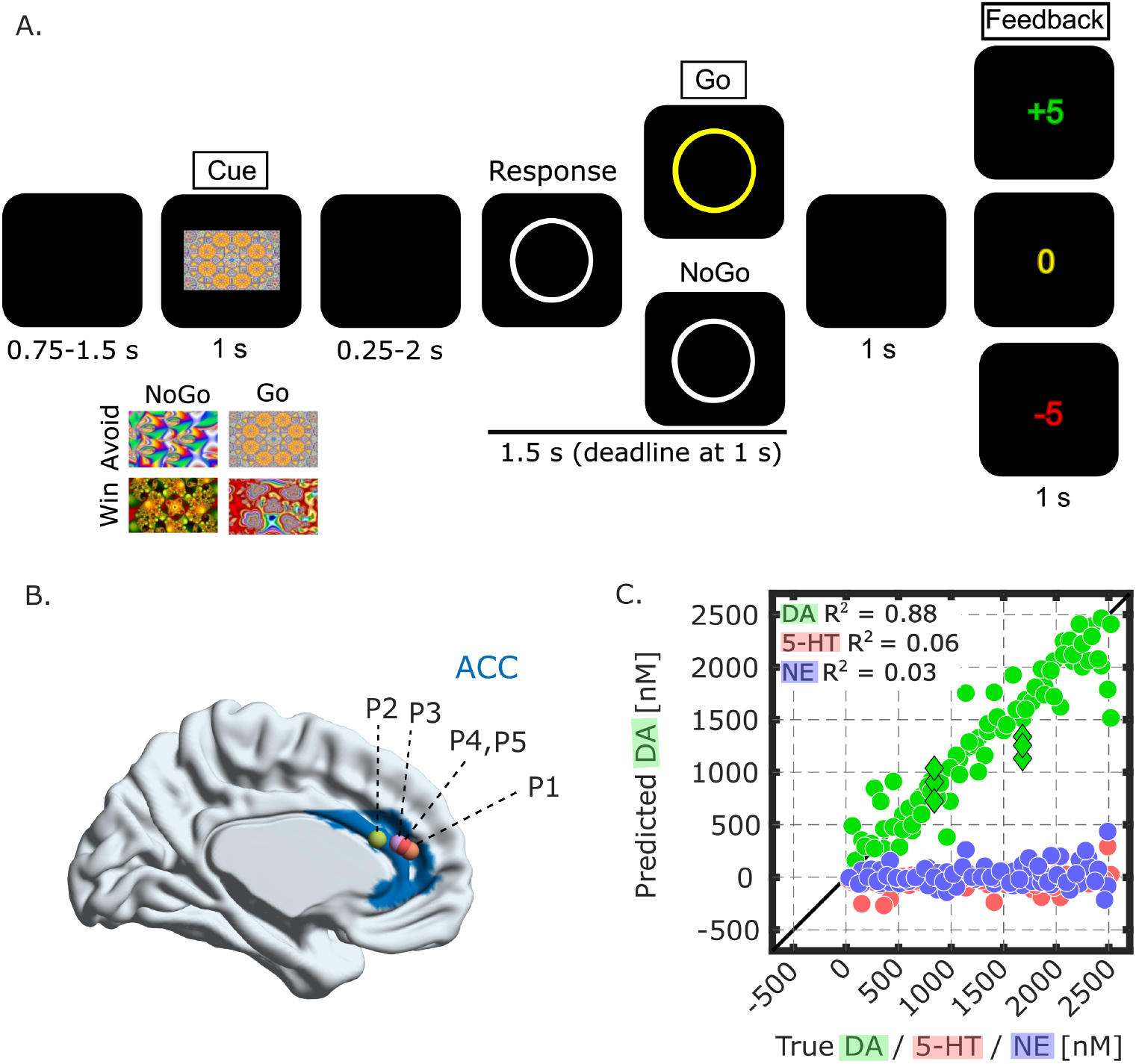
Behavioural task and electrochemical recordings. **A**, Orthogonalised motivational Go/NoGo task. Participants were first presented with one of four cues, with each cue corresponding to a specific state. Participants were then presented with a white circle, at which point they had to decide whether to press (Go) or not press (NoGo) a button. If they pressed within the 1-s deadline, then the circle turned yellow. A correct response led to the better/worse outcome 80%/20% of the time, and vice versa for incorrect responses. In Win states, the possible outcomes were 0 and +5. In Avoid states, the possible outcomes were -5 and 0. Boxed labels indicate neural events of interest. **B**, View of recording sites at midline of brain (X = 0). Dots indicate patients (P) and shaded blue area indicates the ACC. See Methods for coordinates in standard Montreal Neurological Insitute (MNI) space. **C**, In-vitro out-of-training evaluation of electrochemical approach. Dots indicate the average predicted DA concentration (nanomoles, nM) for single-analyte solutions that only contained DA (green), serotonin (5-HT; red), or norepinephrine (NE; blue). Green diamonds indicate mixture solutions that contained 5-HT and/or NE in addition to DA. Predictions were from a 10-fold cross-validation and pooled across patients. R^2^ values were obtained by regressing predicted DA against true DA, 5-HT, or NE. The results show that the approach can detect changes in DA and does not confuse changes in DA for 5-HT or NE.

### 2.2 Behavioural results

To unpack how participants performed the task, we first ran a trial-by-trial logistic mixed-effects model in which we predicted their Go/NoGo responses using the required action in the current state, the valence of the current state and their interaction (response model 1 (R-M1); see Methods for details about statistical analyses). In keeping with earlier studies (Guitart-Masip, Huys, et al., 2012; Moutoussis et al., 2018), participants displayed a Go bias, responding Go on 66% of trials, despite Go only being the required action on 50% of trials (average across states in left part of **Figure 2A** where dark symbols represent patient data; R-M1: intercept, *t*(596) = 3.21, *p* = .001, *β* = 0.72, 95% CI = [0.28, 1.16]).

**Figure 2.**
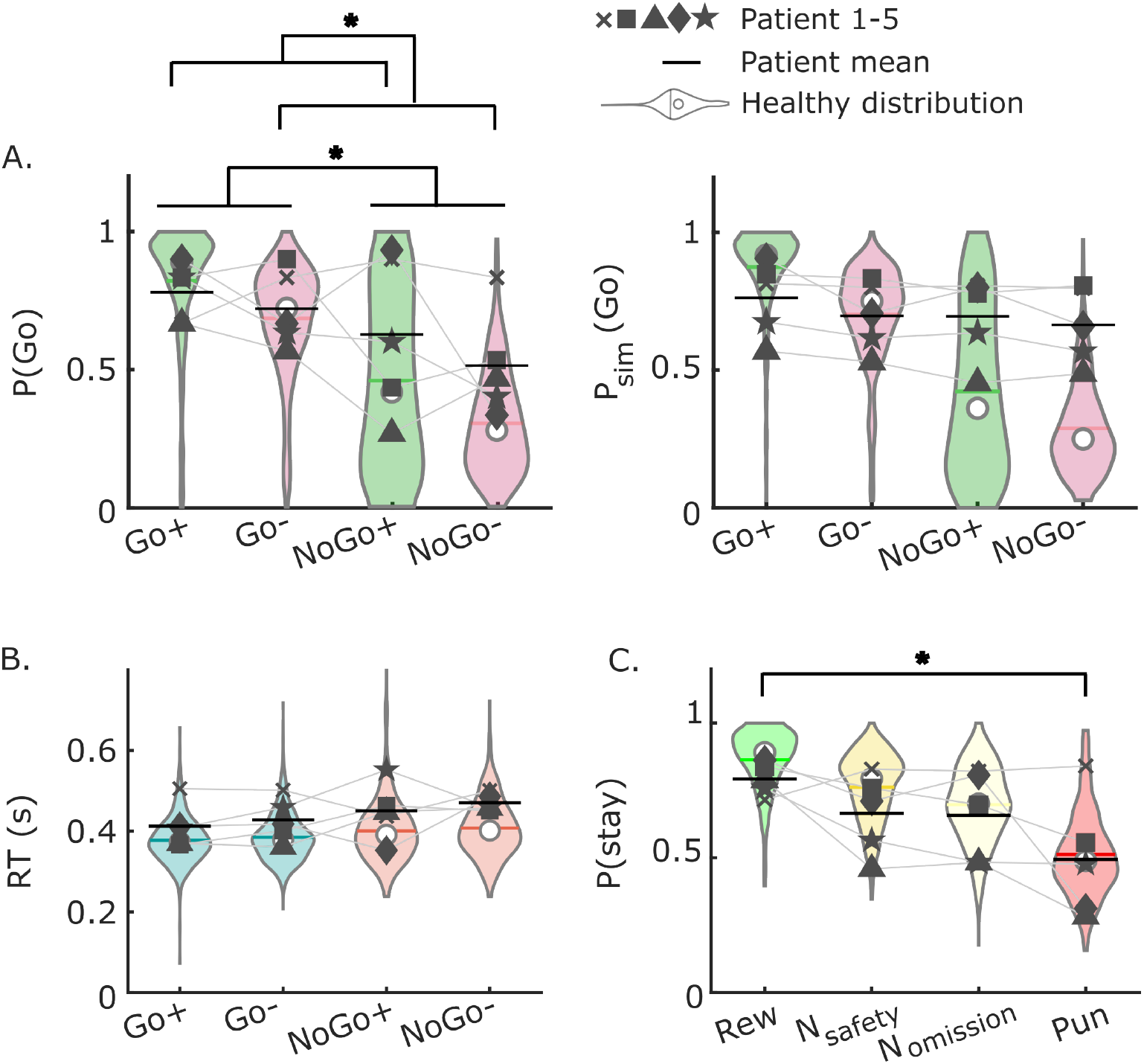
Behavioural results. **A**, Probability of a Go response, P(Go), in each state (left) as observed empirically and (right) as simulated by the reinforcement learning model fitted to the empirical data. For the simulated patient data, each data point was averaged across 100 simulations to account for probabilistic response selection. Green indicates rewarding states, whereas red indicates punishing states. **B**, Reaction time (RT) in seconds (s) in each state. Pale blue denotes RTs for correct Go responses, whereas pale salmon denotes RTs for incorrect Go responses. **C**, Probability of repeating the response made the last time that a state was visited separated by the outcome experienced on that state visit. Rew: reward. N_safety_: neutral outcome in a punishing state. N_omission_: neutral outcome in a rewarding state. Pun: punishment. In **A-C**, dark symbols indicate the patient data, with the black horizontal bar representing the patient mean; the violin plots indicate the distribution of data observed in a cohort of *N* = 817 healthy participants (Moutoussis et al., 2018), with the white dot representing the median and the coloured line representing the mean. Asterisks indicate significant results (*p* < .05) for the patient data.

Participants nevertheless adapted their responses to the task requirements, responding Go on 75% of Go trials compared to 57% of NoGo trials (compare Go and NoGo states in left part of **Figure 2A**; R-M1: state action, *t*(596) = 4.68, *p* < .001, *β* = 0.43, 95% CI = [0.25, 0.60]). In addition, and indicative of Pavlovian biases being in play, participants’ response tendencies were modulated by the valence of a state, with participants responding Go on 70.3% of Win trials compared to 61.7% of Avoid trials, despite the requirement for Go/NoGo being equally balanced within each context (compare Win and Avoid states in left part of **Figure 2A**; R-M1: state valence, *t*(596) = 2.25, *p* = .025, *β* = 0.20, 95% CI = [0.03, 0.38]). The interaction between state action and state valence did not significantly modulate trial-by-trial responses, indicating that the magnitude of the motivational effect of state valence did not vary with the required action (R-M1: *t*(596) = -0.42, *p* = .675, *β* = -0.04, 95% CI = [-0.22, 0.14]).

A linear mixed-effects model of Go reaction times (RTs) using the same predictors did not return significant effects, although there was a trend for correct Go responses being faster than incorrect ones (compare colours in **Figure 2B**; RT model 1 (RT-M1); all absolute *t*(392) < 1.42, all *p* > .153). Critically, despite the unique setting of the EMU, the patients’ behavioural metrics fell within the ranges observed in an earlier study in which a large healthy cohort (*N* = 817) performed the same task (Moutoussis et al., 2018; compare dark symbols to violin plots in **Figure 2**).

Having established general behavioural adaptation to the task requirements, we next tested for trial-by-trial learning effects. Here, we again predicted participants’ Go/NoGo responses in a logistic mixed-effects model, but we now used predictors pertaining to the last time that the current state was visited, which on average was 3.8 trials in the past (R-M2). These historical predictors included the response made on the last state visit and two separate outcome terms that depended on whether the outcome received on the last state visit was unambiguous as well as the interaction between these outcome terms and the last response. More specifically, one of the outcome terms contrasted unambiguous positive and negative outcomes, whereas the other term contrasted ambiguous neutral outcomes on Win trials (omission, as reward is not received) and Avoid trials (safety, as punishment is avoided).

This analysis revealed that, while participants had a general tendency to repeat the response from the last state visit, this tendency was modulated by the unambiguous, although not the ambiguous, outcome history (**Figure 2C**; R-M2: last response, *t*(594) = 5.28, *p* < .001, *β* = 0.50, 95% CI = [0.31, 0.69]; unambiguous outcome, *t*(594) = 0.34, *p* = .731, *β* = 0.25, 95% CI = [-1.16, 1.66]; unambiguous interaction, *t*(594) = 5.02, *p* ¡ .001, *β* = 3.62, 95% CI = [2.20, 5.04]; ambiguous terms, all absolute *t(*594) < 1.14, all *p* > .254). This result suggests that, akin to a win-stay-lose-shift strategy, participants tended to repeat their response on the next state visit after a reward and to change their response after a punishment (**Figure 2C**). Indeed, when running a logistic mixed-effects model with separate outcome terms for unambiguous rewards and punishments instead of one combined unambiguous term as before, we found that rewards increased the tendency to repeat the response from the last state visit, whereas punishments decreased the tendency to repeat the response (R-M2-F1: last response × reward, *t*(594) = 2.35, *p* = .019, *β* = 0.58, 95% CI = [0.10, 1.06]; last response × punishment, *t*(594) = -3.76, *p* < .001, *β* = -0.88, 95% CI = [-1.34, -0.42]).

Taken together, the behavioural results show that, while participants acquired appropriate cue-outcome mappings through instrumental learning, they were subject to Pavlovian biases. From an instrumental viewpoint, they adapted their responses to the required action in a state. From a Pavlovian viewpoint, they also exhibited Pavlovian tendencies to approach (Go) in rewarding contexts (Go+, NoGo+) and avoid (NoGo) in aversive ones (Go-, NoGo-), after controlling for a general Go bias.

### 2.3 Computational modelling

In keeping with earlier work (e.g., Algermissen et al., 2024; Guitart-Masip, Huys, et al., 2012; Swart et al., 2018), we fitted an RL model to participants’ Go/NoGo responses (see Methods for details on modelling). Our aim was to derive predictors for neural analysis that better reflected the cognitive dynamics underpinning task behaviour. We used the RL model which provided the best fit in the large healthy cohort reported in **Figure 2** (Moutoussis et al., 2018).

The model is a variant of Q-learning and includes an instrumental module, which maintains estimates of state-action values (*Q*_*t*_(*s*_*t*_, *a*_*t*_) for state *s*_*t*_ and action *a*_*t*_ on trial *t*), and a Pavlovian module, which only maintains estimates of state values (*V*_*t*_(*s*_*t*_)). The modules update their value estimates using RPEs over state-action and state value expectations, respectively. To capture any asymmetric impact of reward versus punishment, the model has separate learning rates for the two outcomes, with their relative magnitude determining the degree to which state-action and state value estimates are optimistic (reward > punishment) or pessimistic (reward < punishment) (Mihatsch and Neuneier, 2002). The value estimates from the instrumental and Pavlovian modules are combined into action weights for probabilistic (SoftMax) response selection, with a Pavlovian parameter controlling the degree to which the Pavlovian module biases responses towards approach (Go) in rewarding contexts and avoidance (NoGo) in aversive ones. The model also includes a Go bias to capture any general tendency to respond Go regardless of state information.

As shown in the right part of **Figure 2A**, the model, which was fitted at the individual level, broadly captures task behaviour in the patient data (dark symbols) and the large healthy cohort (violin plots). We used the best-fitting parameters for each individual patient dataset to generate model-dependent predictors for neural analysis, specifically trial-by-trial state value estimates, state-action value estimates, and RPEs for the state-action value estimates. We highlight that the model-dependent predictors were generated under the empirically observed responses and that the purpose of using the best-fitted parameters is therefore mainly to scale the model-dependent predictors. Previous work has shown that even large errors in fitted parameters have a minimal impact on the results of model-dependent neural analysis, especially when the model-dependent predictors are z-scored before being used for analysis as here (Wilson and Niv, 2015).

### 2.4 Electrochemical results

We next turned to the electrochemical recordings from the ACC (see Methods for details on neural analysis). We focused on three trial events: the visual cue which signals the current state, the initiation of a Go response, and feedback delivery. In brief, for each event of interest, we created a 10-s epoch for each trial centred on the event onset, z-scored this epoch, and then calculated a trial-level DA estimate as the mean DA estimate across a 1-s window (N = 10 DA estimates) leading up to or starting at the event onset. We used these trial-level DA estimates for statistical analysis, but we visualise results using the time series data. For each event of interest, we report model-agnostic analyses using predictors informed by the task design and model-dependent analyses using predictors from the RL model fitted to task behaviour. Unlike the model-dependent approach, the model-agnostic approach is not constrained, or biased, by our specific choice of model.

To aid interpretation of the electrochemical results, we provide, for each event of interest, cartoons of the DA response expected under standard accounts of DA to accompany the observed DA response. These cartoon models should be treated as abstractions, rather than quantitative predictions. For convenience, they will be simplified to be symmetric between conditions, whereas true predictions are not symmetric. For example, the learned absolute value of negative states should be smaller than that of positive states, because participants will not receive the same amount of punishments as rewards when performing the task at a better than chance level.

#### 2.4.1 Dopamine signals Pavlovian congruence of upcoming actions

We first considered the DA response to the visual cue which signals the current state. We calculated this response at the trial level as the mean DA estimate across a 1-s window starting at cue onset. As illustrated by the cartoon models in the top and middle part of **Figure 3A**, standard expectations would be for DA to increase in anticipation of reward compared to punishment (top; Montague et al., 1996; Schultz, 2025; Schultz et al., 1997; Watabe-Uchida and Amo, 2025). DA should also be greater when required to, or about to, respond Go compared to NoGo (middle; da Silva et al., 2018; Guitart-Masip et al., 2014). To test these hypotheses, we ran a linear mixed-effects model in which we predicted the DA cue response using predictors defined by the task, including the valence of the current state, the required action in the current state and the action about to be performed by the participant (neural model 1 (N-M1)). In light of the behavioural results and earlier animal results (Lloyd and Dayan, 2023; Oleson et al., 2012; Syed et al., 2016), we also allowed for the possibility that DA already at this stage of a trial is modulated by motivational biases, by additionally including interactions between state valence, the required action, and the performed action in the mixed-effects model.

**Figure 3.**
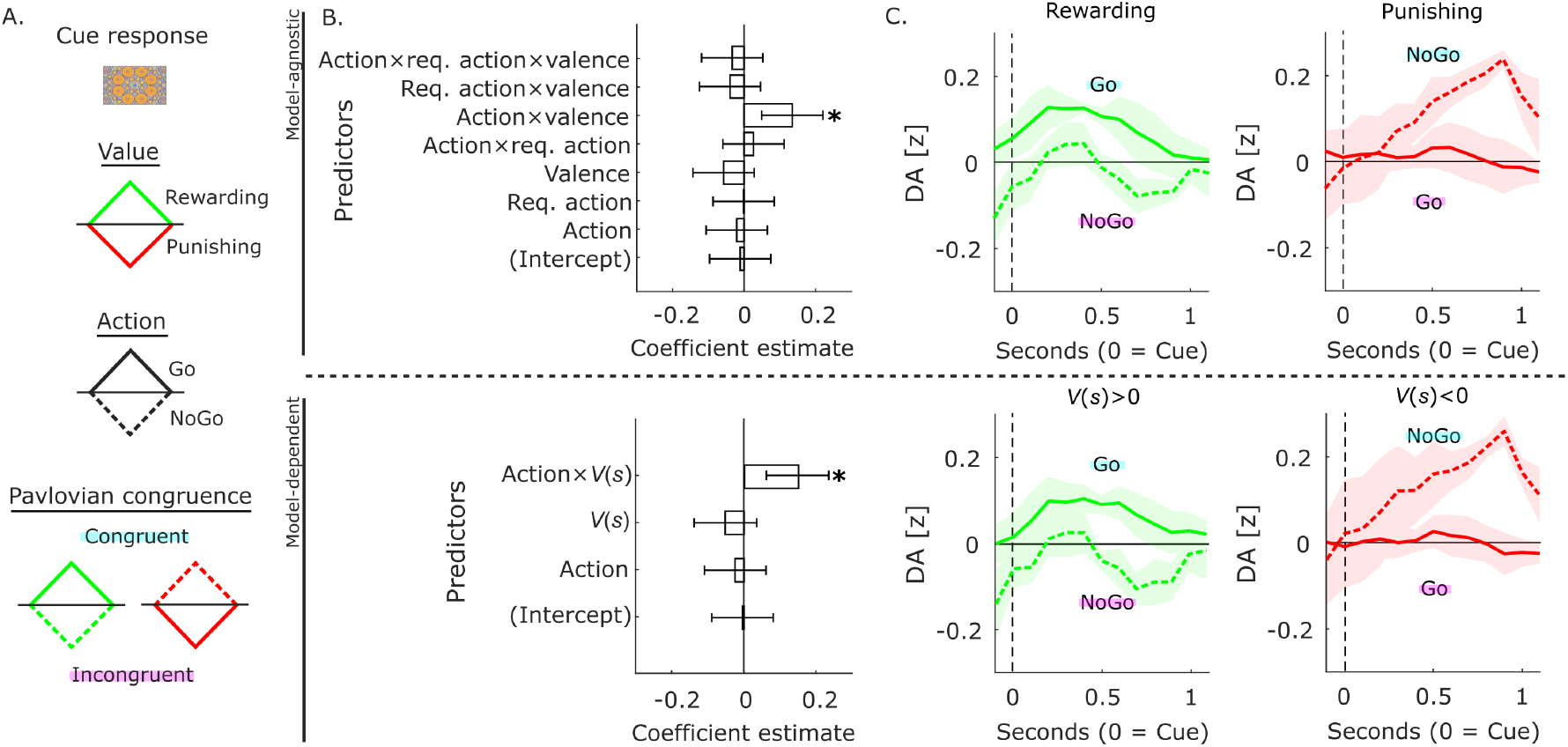
Dopamine signals Pavlovian congruence of upcoming actions. **A**, Schematic of the DA cue response expected under (top) value coding, (middle) action coding and (bottom) modulation by Pavlovian congruence. **B**, Regression results for (top) model-agnostic and (bottom) model-dependent analyses. Asterisks indicate significant results (*p* < .05). **C**, Visualisation of estimated DA time series in (left) rewarding and (right) punishing states separated by the performed action using (top) model-agnostic or (bottom) model-dependent variables. Colours and line styles map onto the cartoon models in **A**. The pattern of responses cannot be fully reconciled with the standard value and action accounts but appears instead related to Pavlovian congruence. In **B-C**, the trial indicator is omitted from the model-dependent predictors and data are represented as the mean ± standard error of the mean across participants.

This model-agnostic analysis revealed a significant interaction between state valence and the performed action (top part of **Figure 3B**; N-M1: action × valence, *t*(592) = 3.10, *p* = .002, *β* = 0.14, 95% CI = [0.05, 0.22]), indicating that DA increases ahead of Pavlovian congruent, relative to Pavlovian incongruent, actions. To unpack this interaction effect, we separated trials into rewarding and punishing states and compared trials with Go and NoGo responses within each context (N-M1-F1; top part of **Figure 3C**). We found that DA levels were higher when the action about to be performed on the current trial was Pavlovian congruent regardless of state valence. Specifically, DA levels were higher for Go than NoGo responses in rewarding states (top-left part of **Figure 3C**; N-M1-F1: *t*(298) = 1.63, *p* = .104, *β* = 0.10, 95% CI = [-0.02, 0.22]) and higher for NoGo than Go responses in punishing states (top-right part of **Figure 3C**; N-M1-F1: *t*(298) = -2.75, *p* = .006, *β* = -0.16, 95% CI = [-0.27, -0.04]). This pattern of responses cannot be fully reconciled with the value and the action cartoon models in **Figure 3A**, as neither account predicts that DA should be higher for NoGo than Go responses in punishing states.

We next ran a linear mixed-effects model in which we predicted the DA cue response using predictors from our learning model which should better capture the cognitive dynamics underpinning task behaviour (N-M2; bottom part of **Figure 3B**). In light of the model-agnostic results, our model-dependent predictors included the learned value of the current state, *V*_*t*_(*s*), the action about to be performed by the participant on the current trial and the interaction between the two terms. In further support of a neural effect of Pavlovian congruence, the interaction between learned state value and performed action was significant (N-M2: state value (*V*_*t*_(*s*)), *t*(596) = -1.16, *p* = .246, *β* = -0.05, 95% CI = [-0.14, 0.03]; action, *t*(596) = -0.55, *p* = .586, *β* = -0.02, 95% CI = [-0.11, 0.06]; interaction, *t*(596) = 3.38, *p* < .001, *β* = 0.15, 95% CI = [0.06, 0.23]). In line with the model-agnostic analysis, DA levels were trending towards being higher for Go than NoGo responses in rewarding states (bottom-left part of **Figure 3C**; N-M2-F1: *t*(258) = 1.66, *p* = .099, *β* = 0.11, 95% CI = [-0.02, 0.24]) and were significantly higher for NoGo than Go responses in punishing states (bottom-right part of **Figure 3C**; N-M2-F1: *t*(298) = -2.75, *p* = .006, *β* = -0.16, 95% CI = [-0.27, -0.04]).

Taken together, these results indicate that neither of the value or action cartoon models in **Figure 3A** is correct. Instead, as illustrated in the bottom part of **Figure 3A**, the results indicate that DA at the level of the ACC signals the Pavlovian congruence of upcoming actions, in line with a role in promoting Pavlovian responding.

#### 2.4.2 Dopamine signals action errors and Pavlovian deviation

We next turned to the DA response during action initiation, an event which is temporally dissociated from cue presentation by the variable interval preceding the response window (**Figure 1A**). We limited our analysis to Go trials (66% of trials) as the task does not provide any precise temporal marker of when a participant chooses NoGo. We calculated the DA action initiation response at the trial level as the mean DA estimate across a 1-s window leading up to the registration of a button press. As illustrated by the cartoon model in the top part of **Figure 4A**, a standard expectation would be for DA to increase in preparation of the execution of a Go response in the context of reward as opposed to punishment (Guitart-Masip et al., 2014), although we note that the earlier studies in rodent ventral striatum found DA to increase for both active approach and active avoidance (Gentry et al., 2016), or only for correctly performed Go responses (Syed et al., 2016). To test these hypotheses, we ran a linear mixed-effects model in which we predicted the DA pre-Go response using state valence as defined by the task, the required action as defined by the task, which specifies whether a Go is correct or not, as well as the interaction between state valence and the required action (N-M3).

**Figure 4.**
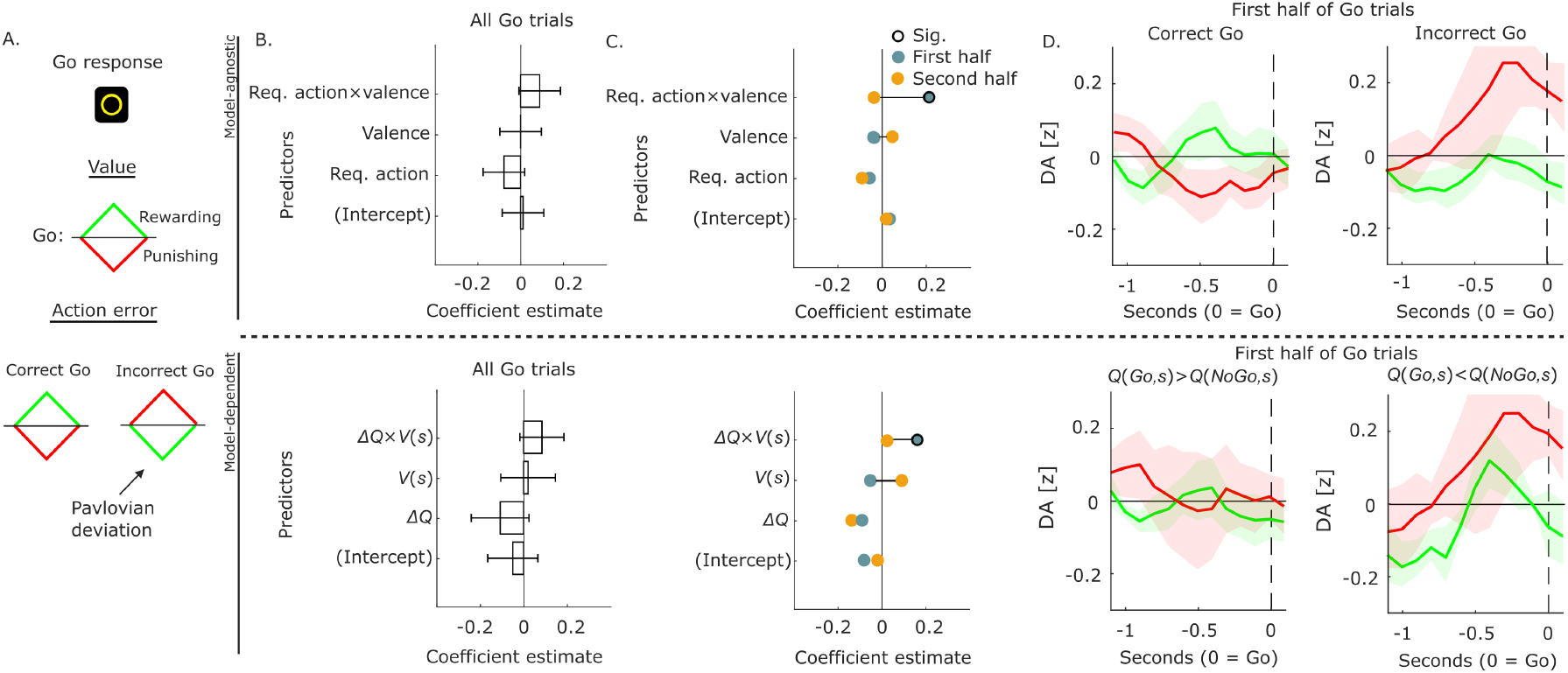
Dopamine signals action errors and Pavlovian deviation. **A**, Schematic of the DA Go response expected under (top) value coding and (bottom) modulation by action errors and deviation from Pavlovian tendencies. **B**, Regression results for (top) model-agnostic and (bottom) model-dependent analyses. Asterisks indicate significant results (*p* < .05). **C**, Same as in **B**, but now for the first (blue) and second (orange) half of Go trials. Sig.: significant (*p* < .05). **D**, Visualisation of estimated DA time series on (left) correct Go trials and (right) incorrect Go trials separated into rewarding and punishing states using (top) model-agnostic or (bottom) model-dependent variables. Colours and line styles map onto the cartoon models in **A**. The pattern of responses cannot be fully reconciled with the standard value account but appears instead related to action errors and Pavlovian deviation. In **B-D**, the trial indicator is omitted from the model-dependent predictors and data are represented as the mean ± standard error of the mean across participants.

This model-agnostic analysis did not return any significant effects (top part of **Figure 4B**; N-M3: all other absolute *t*(392) < 1.82, all *p* > .069). One reason may be that the effects of the task-defined predictors vary across time. For example, the effect of Go errors may be greater later on in the task once task performance has stabilised, or early on when uncertainty about which action to take is greater. To test this possibility, we ran an exploratory analysis in which we separated Go trials into the first and second half of Go trials and repeated the model-agnostic analysis for each half (N-M3; top part of **Figure 4C**). This analysis identified an interaction between state valence and the required action in the first half of Go trials (orange in top part of **Figure 4C**; N-M3, first half: valence, *t*(197) = -0.63, *p* = .531, *β* = -0.04, 95% CI = [-0.18, 0.09]; required action, *t*(197) = -0.89, *p* = .375, *β* = -0.06, 95% CI = [-0.20, 0.07]; interaction, *t*(197) = 3.22, *p* = .001, *β* = 0.22, 95% CI = [0.09, 0.36]), but not in the second half of Go trials (blue in top part of **Figure 4C**; N-M3, second half: all absolute *t*(191) < 1.23, all *p* > .219). To unpack the interaction effect in the first half of Go trials, we separated the trials into correct and incorrect Go responses and compared trials in rewarding and punishing states within each category (N-M3-F1; top part of **Figure 4D**). Consistent with the value cartoon model in **Figure 4A**, during the initiation of a correct Go, DA levels were trending towards being higher in rewarding than punishing states (top-left part of **Figure 4D**; N-M3-F1: *t*(105) = 1.64, *p* = .104, *β* = 0.15, 95% CI = [-0.03, 0.33]). However, in contrast to the value cartoon model, during the initiation of an incorrect Go, DA levels were significantly higher in punishing than rewarding states (top-right part of **Figure 4D**; N-M3-F1: *t*(92) = -2.82, *p* = .006, *β* = -0.27, 95% CI = [-0.46, -0.08]). Notably, from the perspective of Pavlovian tendencies, an incorrect Go in punishing states is more surprising than an incorrect Go in a rewarding state.

For the model-dependent analysis, we used the learned state value of the current state, *V*_*t*_(*s*), the difference between the learned state-action values of Go and NoGo, Δ*Q*_*t*_ = *Q*_*t*_(*Go, s*) − *Q*_*t*_(*NoGo, s*), and their interaction as predictors (N-M4). This analysis also did not return any significant effects (bottom part of **Figure 4B**; N-M4: all other absolute *t*(392) < 1.63, all *p* > .104). However, in line with the model-agnostic analysis, we found an interaction between the learned state value and the learned state-action value difference in the first half of Go trials (blue in bottom part of **Figure 4C**; N-M4, first half: *V*_*t*_(*s*), *t*(197) = -0.64, *p* = .524, *β* = -0.05, 95% CI = [-0.21, 0.11]; Δ*Q*_*t*_, *t*(197) = -0.94, *p* = .349, *β* = -0.09, 95% CI = [-0.27, 0.10]; interaction, *t*(197) = 2.39, *p* = .018, *β* = 0.15, 95% CI = [0.03, 0.28]), but not in the second half of Go trials (orange in bottom part of **Figure 4C**; N-M3, second half: all absolute *t*(191) < 1.43, all *p* > .155). Again, the follow-up analysis showed that, while there was no significant difference when the learned state-action value was higher for Go than NoGo (bottom-left part of **Figure 4D**; N-M4-F1: *t*(87) = 0.78, *p* = .440, *β* = 0.08, 95% CI = [-0.12, 0.29], DA levels were higher for Go in punishing states compared with rewarding ones when the learned state-action value in fact favoured NoGo over Go (bottom-right part of **Figure 4D**; N-M4-F1: *t*(62) = -2.71, *p* = .009, *β* = -0.32, 95% CI = [-0.55, -0.08]).

Taken together, as illustrated in the bottom part of **Figure 4A**, these results indicate that DA at the level of the ACC signals erroneously initiated actions, specifically when a Go error deviates from Pavlovian tendencies. This response may reflect surprise about the initiated action, in line with the recently reported role of DA in signalling action prediction errors, defined as the difference between the action performed and the action expected in the current state given past behaviours (Greenstreet et al., 2025). Alternatively, in keeping with the reported DA cue response, it may reflect a control signal that promotes Pavlovian responding following the detection of a Pavlovian incongruent error that is about to be committed.

#### 2.4.3 Dopamine signalling of reward prediction errors depends on Pavlovian congruence

Finally, we examined the DA response to the trial outcome, which we calculated at the trial level as the mean DA level across a 1-s window starting at feedback delivery. As illustrated by the cartoon model in top part of **Figure 5A**, a standard expectation under the RPE theory of DA (Montague et al., 1996; Schultz et al., 1997) would be for DA to increase when an outcome is better than expected as compared to when worse than expected (top part of **Figure 5A**). To test this hypothesis, we ran a linear mixed-effects model in which we predicted the DA outcome response using the experienced trial outcome, where the trial outcome term separates neutral outcomes in rewarding and punishment states (−1 = punishment, −0.5 = neutral outcome in a rewarding state, 0.5 = neutral outcome in a punishing state and 1 = reward) (N-M5). In light of the earlier results, we also allowed for the possibility that the DA outcome response is modulated by motivational biases, by additionally including the valence of the current state, the required action in the current state as well as the interactions between these terms in the mixed-effects model. Here, the interaction between state valence, the required action, and the outcome, which is affected by the actual action, together determine whether future Pavlovian responding is favoured.

**Figure 5.**
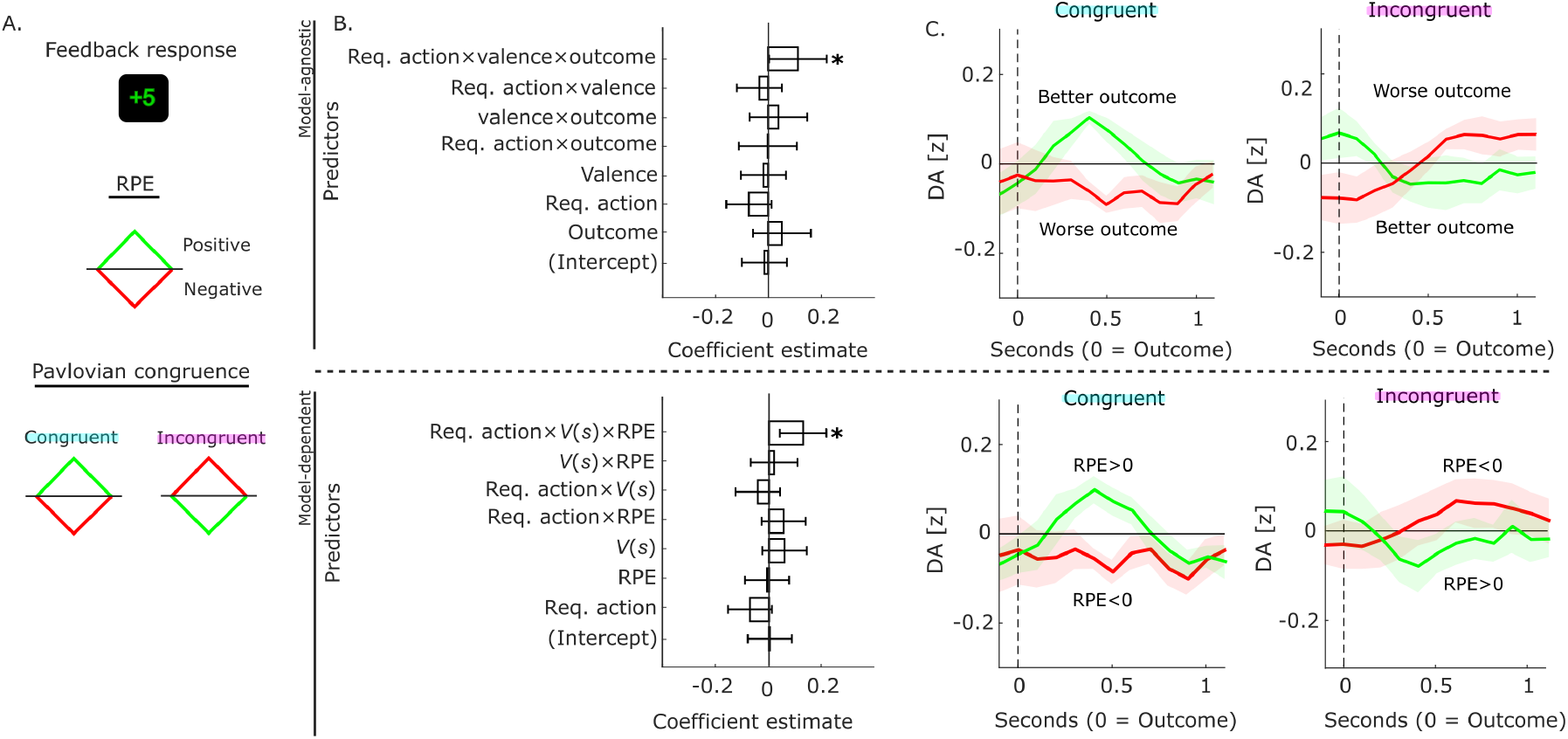
Dopamine signalling of reward prediction errors depend on Pavlovian congruence. **A**, Schematic of the DA cue response expected under (top) the classic RPE theory and (bottom) RPE signalling modulated by Pavlovian congruence. **B**, Regression results for (top) model-agnostic and (bottom) model-dependent analyses. Asterisks indicate significant results (*p* < .05). **C**, Visualisation of estimated DA time series in (left) Pavlovian-congruent and (right) Pavlovian incongruent states separated by the sign of the outcome using (top) model-agnostic or (bottom) model-dependent variables. Colours and line styles map onto the cartoon models in **A**. The pattern of responses cannot be fully reconciled with the standard RPE account but appears instead related to an RPE-like signal promoting Pavlovian responding. In **B-C**, the trial indicator is omitted from the model-dependent predictors and data are represented as the mean ± standard error of the mean across participants.

In line with a role of DA in this process, the analysis revealed a significant interaction between the required action in the current state, the valence of the current state, and the trial outcome (top part of **Figure 5B**; N-M5: req. action × valence × outcome, *t*(592) = 2.01, *p* = .045, *β* = 0.11, 95% CI = [0.00, 0.22]; outcome,, *t*(592) = 0.93, *p* = .354, *β* = 0.05, 95% CI = [-0.06, 0.16]; all other absolute *t*(592) < 1.69, all other *p* > .092). To unpack this interaction effect, we separated the trials according to the Pavlovian congruence of the current state and regressed the outcome against the DA outcome response within each category (N-M5-F1; top-part of **Figure 5C**). While the main effect of outcome was not individually significant within either category, the response pattern indicated that the interaction between Pavlovian congruence and outcome was driven by DA scaling positively with outcome in Pavlovian-congruent states (top-left part of **Figure 5C**; N-M5-F1: *t*(298) = 1.67, *p* = .096, *β* = 0.11, 95% CI = [-0.02, 0.25]) but negatively in Pavlovian-incongruent states (top-right part of **Figure 5C**; N-M5-F1: *t*(298) = -0.65, *p* = .519, *β* = -0.05, 95% CI = [-0.20, 0.10]). While the former result can be reconciled with the RPE cartoon model in **Figure 5A**, the latter result cannot.

For the model-dependent analysis, we replaced state valence as defined by the task with the graded estimate of state value (*V*_*t*_(*s*)) and the trial outcome with the RPE for the state-action value estimate (N-M6; bottom part of **Figure 5B**). In line with the model-agnostic results, this analysis identified an interaction between the required action, state value and RPEs, where the required action and state value together define Pavlovian congruence (bottom part of **Figure 5B**; N-M6: req. action × *V*_*t*_(*s*) × RPE, *t*(592) = 2.87, *p* = .004, *β* = 0.13, 95% CI = [0.04, 0.21]; all other absolute *t*(592) < 1.72, all other *p* > .085). While the main effect of RPE was not individually significant when separately analysing Pavlovian-congruent and Pavlovian-incongruent states (N-M6-F1; bottom part of **Figure 5C**), the response pattern again indicated that the interaction between Pavlovian congruence and RPE was driven by DA scaling positively with RPEs in Pavlovian-congruent states (bottom-left part of **Figure 5C**; N-M6-F1: *t*(272) = 1.83, *p* = .069, *β* = 0.11, 95% CI = [-0.00, 0.23]) but negatively in Pavlovian-incongruent states (bottom-right part of **Figure 5C**; N-M6-F1: *t*(253) = -1.01, *p* = .313, *β* = -0.06, 95% CI = [-0.18, 0.06]).

Taken together, as illustrated in the bottom part of **Figure 5A**, these results indicate that DA at the level of the ACC does not signal a standard RPE but instead an RPE-like signal that promotes future Pavlovian responding. As observed, if Pavlovian responding is the default policy being reinforced, such a signal should be higher not only for better than expected outcomes in Pavlovian-congruent states but also higher for worse than expected outcomes in Pavlovian-incongruent states, with the latter scenario implying that Pavlovian responding would have been favourable.

## 3 Discussion

In this study, we investigated how Pavlovian-instrumental conflict engages the DA system, by performing intracranial electrochemical recordings of fast DA dynamics in human ACC during a motivational Go/NoGo task. The task fully orthogonalises the required action in a state (Go versus NoGo) and the valance of a state (reward versus punishment). It involves two Pavlovian congruent states, where instrumental and Pavlovian responding favour the same action (Go-to-Win and NoGo-to-Avoid), and two Pavlovian incongruent states, where approach bias in rewarding contexts and avoidance bias in aversive ones directly conflict with instrumental action selection (Go-to-Avoid and NoGo-to-Win) (Crockett et al., 2009; Guitart-Masip, Huys, et al., 2012; Guitart-Masip et al., 2014). We did not find the standard motor or value signals, such as canonical RPEs, typically seen in striatal DA recordings (da Silva et al., 2018; Guitart-Masip et al., 2014; Schultz, 2025; Schultz et al., 1997; Watabe-Uchida and Amo, 2025). Instead, we found, in both model-agnostic analyses and analyses based on an RL model fitted to task behaviour, that these signals were coded in the frame of reference of Pavlovian biases. Our results indicate that DA at the level of the ACC is involved in evaluating the degree to which a default policy — in this case Pavlovian responding — should guide behaviour.

We observed this computational motif across three key task stages: during the presentation of the visual cue which signals the state on the current trial; during the initiation of Go responses; and during the presentation of the outcome received on the current trial. Immediately following cue onset, we observed an increase in DA specifically when participants later in that trial performed a Pavlovian-congruent action (Go-to-Win and NoGo-to-Avoid) rather than a Pavlovian-incongruent action (Go-to-Avoid and NoGo-to-Win) (**Figure 3**). These DA dynamics diverge from, or refine, the pattern seen in striatal recordings in animals. For example, one study reported cue-evoked increases in DA in rat ventral striatum ahead of active approach for reward (Go-to-Win) and avoidance of punishment (Go-to-Avoid) as compared to neutral settings (Gentry et al., 2016). In the same species and region, another study reported cue-evoked increases in DA ahead of correctly performed approach for reward (Go-to-Win) as compared to when approach was not needed (NoGo-to-Win) (Syed et al., 2016). Rather than signalling the motivational salience of cues and activating behaviour, our results indicate that DA at level of ACC instead signals Pavlovian salience and promotes Pavlovian responding.

During the initiation of Go responses, we observed an increase in DA when a Pavlovian-incongruent error was imminent, specifically when the participant was about to press a button when the trial required them to refrain from pressing in order to avoid a punishment (NoGo-to-Avoid) (**Figure 4**). This response pattern is in line with the well-documented role of the ACC in error processing. The ACC is believed to be the source of the error-related negativity (ERN) often observed in human scalp EEG (Gehring et al., 2018; Yeung et al., 2004), and it has been hypothesised that the ERN is generated when the DA system transmits negative RL-like signals to the ACC and that the ACC uses these signals to regulate task performance (Holroyd and Coles, 2002). Our results indicate that, rather than signalling just any type of error, the DA system instead reports whether the error is a deviation from a default behavioural policy, here Pavlovian responding. The response pattern is also consistent with the reported role of DA in signalling action prediction errors, defined as the difference between the action performed and the action expected in the current state given behavioural tendencies (Greenstreet et al., 2025). However, unlike the standard ERN or action prediction errors, we observed the increase in DA immediately before and not after an error. This timing indicates that, rather than an error signal per se, the DA system may transmit a correction signal in a final attempt to promote Pavlovian responding.

Immediately following feedback delivery, we found that the DA outcome response depended on whether the outcome favoured Pavlovian responding (**Figure 5**). In Pavlovian-congruent states (Go-to-Win and NoGo-to-Avoid), we observed canonical RPE signalling, with an increase in DA for outcomes that were better than expected relative to worse than expected. In contrast, in Pavlovian-incongruent states (Go-to-Avoid and NoGo-to-Win), this RPE signalling flipped, with an increase in DA for outcomes that were worse than expected relative to better than expected. Rather than simply updating value estimates for states and actions as often reported in the striatum (Schultz, 2025; Schultz et al., 1997; Watabe-Uchida and Amo, 2025), these results indicate that DA at the level of the ACC is involved in updating estimates of the value of a default behavioural policy. In the context of Pavlovian responding, a learning signal favouring Pavlovian responding should be greater not only after a better-than-expected outcome in a Pavlovian-congruent state as this outcome (usually) follows a Pavlovian-congruent action but also after a worse-than-expected outcome in a Pavlovian-incongruent state as this outcome (usually) follows a Pavlovian-incongruent action. More broadly, this interpretation fits with a proposal that the reliance on instrumental versus Pavlovian responding is dynamically adapted to the environment, with a greater reliance on Pavlovian responding in environments that are less controllable from an instrumental perspective (Dorfman and Gershman, 2019). Although our task did not manipulate controllability, our results indicate that DA at the level of the ACC may play a critical role in this arbitration process.

In summary, our results indicate that DA at the level of the ACC is involved in a more abstract, policy-level computation where states and actions are not evaluated in isolation, but instead in the frame of reference of a default behavioural policy. In the same way that DA in the ventral striatum appears, sometimes paradoxically, to couple reward and action, DA in the ACC seems, also sometimes paradoxically, to couple reward and Pavlovian/instrumental congruence. Our results cannot be explained by standard motor or habit-based accounts, where the role of DA is to activate behaviour or update action repetition frequencies, or the canonical RPE account, where DA signals whether cues or outcomes are better than expected as computed by standard RL. Instead, our results indicate that DA promotes Pavlovian responding both when behavioural activation and inhibition are required, tracks whether actions deviate from a reflexive heuristic of Pavlovian responding, and signals whether an outcome favours Pavlovian responding. These results, which differ from observations in the striatum, are consistent with the role of the ACC in adaptive behavioural control, including the adaptation of behavioural strategies to environmental demands.

## 4 Methods

### 4.1 Participants

Four participants with epilepsy (2 female, age range: 26-28) took part in the study during their stay in an epilepsy monitoring unit (EMU). One of the participants took part in two study sessions, providing a total of five datasets. The patients had undergone the implantation of depth electrodes in multiple brain regions, including the anterior cingulate cortex (ACC), for the localization of epileptic foci. Prior to the stay in the EMU, participation in the study was discussed with the patient and the clinical team. The study protocol was described verbally and in a written format before the patient provided informed consent. Patients continued taking their regular medications during the study. No adverse or unanticipated events occurred during or as a result of the study. The study was approved by the IRB committee at the University of Arizona (STUDY00000295).

### 4.2 Behavioural task

Participants performed a motivational Go/NoGo task (adapted from Guitart-Masip, Huys, et al., 2012). On each trial, one of four visual cues (fractals) was first presented for 1 s. Each cue corresponded to a specific task state which specified the best possible outcome and which action was most likely to produce this outcome. The states were: Go-to-Win (Go+), Go-to-Avoid (Go-), NoGo-to-Win (NoGo+), and NoGo-to-Avoid (NoGo-). In Win states, the outcome was either a reward (5 points) or neutral (0 points). In Avoid states, the outcome was either neutral (0 points) or a punishment (-5 points). The outcomes were probabilistic, with the correct (incorrect) action leading to the better outcome with 80% (20%) probability and the worse outcome with 20% (80%) probability. Participants had to learn which cue corresponded to which state through trial-and-error.

After a variable post-cue interval (fixation cross for 0.25-2 s), a white circle was shown. Participants had 1 s to decide whether to press a button (Go) or refrain from pressing a button (NoGo). The circle was shown for 1.5 s. If they pressed a button before the 1 s deadline, then the circle turned yellow for the remaining time; if they pressed a button after the deadline, then the circle remained white. Following a fixed post-action interval (fixation cross for 1 s), participants were presented with feedback for 1 s: a green upward arrow indicated a reward (+5 points); a red downward arrow indicated a punishment (-5 points); and a yellow horizontal bar indicated a neutral outcome (0 points). Finally, there was a variable inter-trial interval (blank screen for 0.75-1.5 s) before participants continued to the next trial. There were a total of 120 trials (40 trials per cue).

### 4.3 Electrochemical recordings

The approach for obtaining sub-second electrochemical estimates of DA from clinical depth electrodes in epilepsy patients has been described in detail in previous work (Bang et al., 2023; Batten et al., 2025). In brief, the neurosurgeon implants the depth electrodes into potential epileptic loci. Here, the electrodes were Ad-Tech Behnke-Fried electrodes (WB09R-SP00X-0B6). During their subsequent stay in the EMU, patients who volunteer to participate in research perform a behavioural task while an electrode in a region of interest is used for fast-scan cyclic voltammetry (FSCV).

Here, we recorded from the ACC, using micro-wire 9 as the reference electrode and one of the remaining wires as the working electrode. The current response at the working electrode is sampled at 100 KHz while a standard triangular voltage waveform is applied at 10 Hz. We specified the waveform as follows: ramp up from –0.6 V to +0.175 V at 155 V/s, ramp down from +0.175 V to –0.6 V at –155 V/s, hold at –0.6 V for 90 ms. Once epilepsy monitoring is complete, the neurosurgeon explants the electrodes, and the electrode(s) used for research is transported to the laboratory. An electrode-specific signal prediction model for electrochemical estimation is trained by exposing the relevant micro-wire-pairing to labelled concentrations of DA, serotonin (5-HT), norepinephrine (NE) and pH in a controlled in-vitro setting. **Figure 1C** shows results from an in-vitro model evaluation where each electrode-specific model is applied to data withheld from model training. Finally, to generate in-vivo electrochemical estimates, the model is applied to the FSCV data recorded during task behaviour.

### 4.4 Recording sites

The electrodes were reconstructed in patient space and then mapped onto standard Montreal Neurological Institute (MNI) space using the Lead-DBS toolbox (version 3.2.1; Horn and Kühn, 2015; Horn et al., 2019) in MATLAB (2024b, MathWorks). Using the dataset labelling in **Figure 1B**, the MNI coordinates of the recording sites were: P1 = [X, Y, X] = [-1, 42, 15]; P2 = [0, 28, 20]; P3 = [-0.5, 37, 18]; and P4/P5 = [0, 39, 17]. The visualisation in **Figure 1B** was created using the BrainNet Viewer toolbox (version 1.7; Xia et al., 2013) in MATLAB (2024b, MathWorks).

### 4.5 Computational modelling

We modelled task behaviour using the reinforcement learning (RL) model that provided the best fit for the large healthy cohort reported reported in **Figure 2** (Moutoussis et al., 2018). The model, which is a variant of Q-learning, contains an instrumental module, which promotes actions based on state-action-outcome associations, and a Pavlovian module, which promotes Go in states associated with reward and NoGo in states associated with punishment.

The instrumental module maintains estimates of state-action values, *Q*(*s, a*), where *s* indicates the state (cue) and *a* indicates the action (Go or NoGo). The module uses a simple Rescorla-Wagner rule to update the estimate of the value of performing action *a* in state *s* based on what happens on a trial:

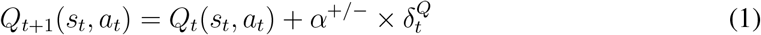

Here, *t* indicates the trial number, *s*_*t*_ indicates the state on that trial, *a*_*t*_ indicates the action performed on that trial, *α*^+/−^ is a learning rate, which can be different for positive and negative outcomes, and 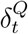 is a reward prediction error (RPE) for the state-action value estimate:

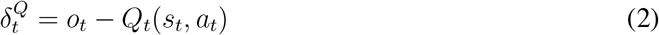

Here, *o*_*t*_ is the outcome (punishment = -5, neutral = 0, reward = 5).

The Pavlovian module maintains estimates of state values, *V* (*s*), and updates these estimates using the same Rescorla-Wagner rule:

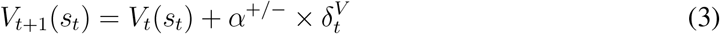

The learning rate *α*^+/−^ is shared with the instrumental module and 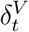 is an RPE for the state value estimate:

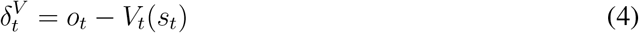

The models combine the instrumental and Pavlovian modules to generate action propensities, *W*_*t*_(*s*_*t*_, *a*), that determine the probability of choosing action *a* on trial *t*:

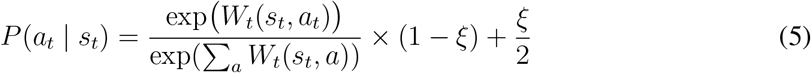

Here, *ξ* is a lapse parameter that accounts for irreducible noise in the response process. The action propensities are defined as:

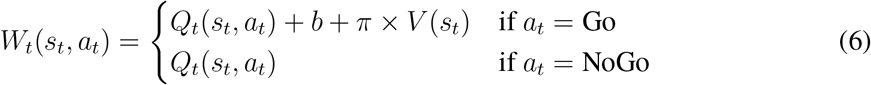

Here, *b* is a Go bias, which captures any general propensity to respond Go, and π is a Pavlovian bias, which controls the degree to which the learned value of a state promotes, or suppresses, Go relative to NoGo.

We fitted the model separately for each dataset, using the dataset-level fitting procedure implemented in the Computational and Behavioral Modeling Toolbox (Piray et al., 2019), which uses MATLAB’s (R2024b, MathWorks) fminunc function to fit parameters, by maximizing the model likelihood given the behavioural data. We used the best-fitting dataset-level parameters and behavioural datasets as input to the model to compute trial-by-trial changes in model variables, like state value estimates (*V*_*t*_(*s*)), state-action value estimates (*Q*_*t*_(*s, a*)) and RPEs for the state-action value estimates. These variables then used as model-dependent predictors for neural analysis.

### 4.6 Statistical analysis

#### 4.6.1 Mixed-effects models

We used mixed-effects models specified at the trial level for statistical analysis of both behavioural and neural data. All models included fixed (group-level) effects and a random intercept varying by dataset. We note that, for all reported analyses, a model which only included a random intercept provided a better fit (AIC/BIC) than a model which also included random effects. All statistical tests were two-tailed. The mixed-effects models were implemented using the ‘fitglme’ function in MATLAB (R2024b, MathWorks). We use Wilkinson notation to describe the models. We omit the trial indicator, *t*, for simplicity.

#### 4.6.2 Behavioural analysis

We used logistic mixed-effects models to analyse participants’ responses (R; NoGo = 0, Go = 1).

We first ran the following model:

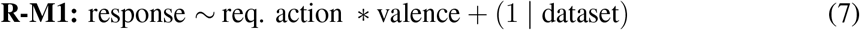

In this model, the required action (NoGo = -1, Go = 1) and the valence (punishment = -1, reward = 1) of the current state are dictated by the task.

To test for trial-by-trial learning effects, we ran the following model:

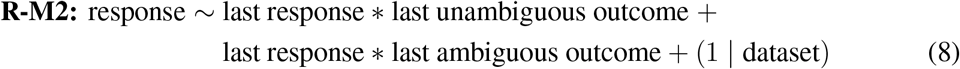

In this model, the historical terms are for the response made (NoGo = -1, Go = 1) and the outcome received the last time that the current state was visited. One outcome terms contrasts unambiguous outcomes (punishment = -1, neutral = 0, reward = 1) and the other outcome term contrasts ambiguous outcomes (omission = -1, reward/punishment = 0, safety = 1).

In a follow-up analysis, we ran the following model:

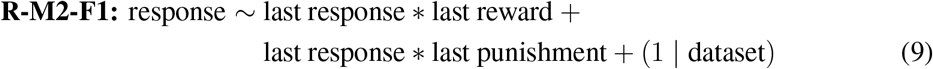

In this model, the unambiguous outcomes were coded as 1 if present and otherwise as 0.

We used a linear mixed-effects model to analyse Go reaction times (RTs), which were first log-transformed and then z-scored. The model was specified as follows:

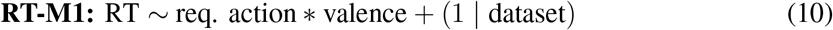

#### 4.6.3 Neural analysis

We first epoched the DA time series using our task events of interest: cue presentation, action initiation (Go trials only), and feedback delivery. Each epoch spanned a 10 s window from 5 s before until 5 s after the task event. We first linearly detrended each epoch to correct for any signal drift and then z-scored each epoch to correct for any temporal variation in signal amplitude. These steps reduce the influence of technical artifacts and facilitate both within-subject and group-level analysis by focusing on relative DA fluctuations, and they have previously been shown to reveal task-related changes in neuromodulator dynamics (e.g., (Bang et al., 2020; Batten et al., 2025). Finally, for statistical analysis, we computed single-trial DA estimates as the area under the curve within a 1-s window leading up to (initiation) or starting at (cue and feedback) the task event.

We describe below the linear mixed-effects models used for neural analysis. All continuos variables, including single-trial DA estimates and model-dependent predictors, were z-scored at the level of datasets. However, in follow-up analyses that focus on particular subsets of trials, the continuos predictors were z-scored at the level of each subset.

The model-agnostic analysis of the DA cue response was specified as follows:

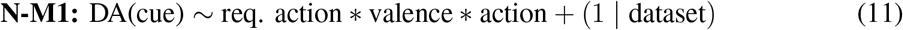

In this model, the required action (NoGo = -1, Go = 1) and the valence (punishment = -1, reward = 1) of the current state are dictated by the task, whereas the last term specifies the action that the participant later performed on the current trial (NoGo = -1, Go = 1). In a follow-up analysis, we ran the following model separately for trials in rewarding (valence > 0) and punishing (valence < 0) states as defined by the task:

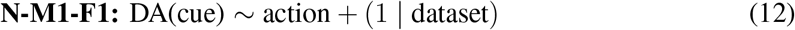

Our model-dependent analysis of the DA cue response was specified as follows:

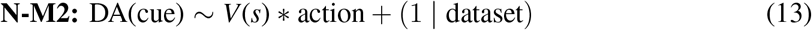

In a follow-up analysis, we ran the following model separately for trials in rewarding (*V* (*s*) > 0) and punishing (*V* (*s*) < 0) states as estimated by the learning model:

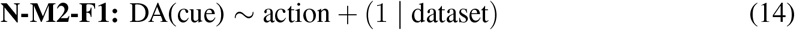

The model-agnostic analysis of the DA action initiation response was specified as follows, both when analysing all Go trials together and when separating Go trials into a first and second half:

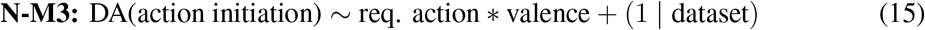

In a follow-up analysis, we ran the following model separately for correct Go trials (req. action > 0) and incorrect Go trials (req. action < 0):

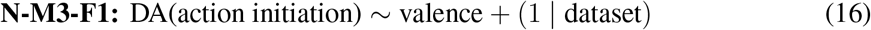

The model-dependent analysis of the DA action initiation response was specified as follows, both when analysing all Go trials together and when separating Go trials into a first and second half:

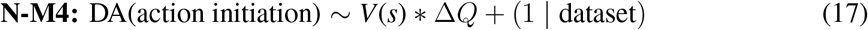

In this model, the second term is the difference in the state-action value estimate between Go and NoGo, Δ*Q* = *Q*(*Go, s*) − *Q*(*NoGo, s*). In a follow-up analysis, we ran the following model separately for Go trials where the state-action value estimate favoured a Go response (Δ*Q* > 0) and Go trials where a NoGo response was favoured (Δ*Q <* 0):

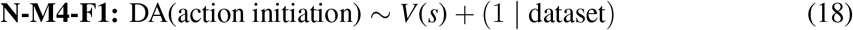

The model-agnostic analysis of the DA outcome response was specified as follows:

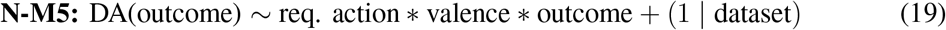

In this model, the outcome term separate both rewards and punishments as well as neutral outcomes indicating omission and safety (punishment = -1, omission = -0.5, safety = 0.5, reward = 1). In a follow-up analysis, we ran the following model separately for Pavlovian-congruent trials (req. action × valence > 0) and Pavlovian-incongruent trials (req. action × valence < 0):

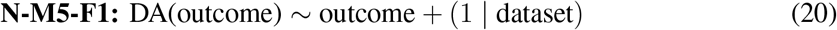

The model-dependent analysis of the DA outcome response was specified as follows:

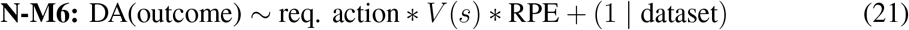

In this model, the RPE was for the state-action value estimate, *δ*^*Q*^. In a follow-up analysis, we ran the following model separately for Pavlovian-congruent trials (req. action × *V* (*s*) > 0) and Pavlovian-incongruent trials (req. action × *V* (*s*) < 0):

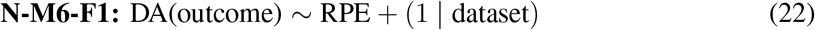

## 5 Data and code availability

Data and code for reproducing figures and associated analyses are available on GitHub: https://github.com/Azadeh-Nazemorroaya/DA-ACC-GNG/.

## 6 Acknowledgements

This work was supported by the Red Gates Foundation (P.R.M.); the National Institute of Mental Health (P.C. and P.R.M.: R01MH122512; G.A.B., S.M.M., R.W.B., and P.R.M.: R01MH132635; P.R.M.: R01MH122948); the Max Planck Society (A.N. and P.D.); the Humboldt Foundation (P.D.); and the Lundbeck Foundation (D.B.: R368-2021-325).

## 7 Author contributions

P.R.M., P.D., and D.B. conceived the study. S.R.B., I.G., A.T., X.C., O.M., C.L., and A.W. collected the data. R.W.B. performed electrode implantation and explantation. D.N. and R.W.B. performed the reconstruction of electrode coordinates. A.N., S.R.B., L.S.B., T.L., and D.B. processed the data. A.N. analysed, modelled, and visualised the data under the supervision of P.D. and D.B. A.N., P.R.M., P.D., and D.B. interpreted the results. A.N. wrote the manuscript under the supervision of P.D. and D.B. The manuscript has been edited by S.R.B., T.L., G.A.B., S.M.M. and P.R.M.. The manuscript has been reviewed by all authors.

